# Development of molecular markers of the Kunitz trypsin inhibitor mutant alleles generated by CRISPR/Cas9-mediated mutagenesis in soybean

**DOI:** 10.1101/2022.08.22.504807

**Authors:** Zhibo Wang, Zachary Shea, Luciana Rosso, Chao Shang, Jianyong Li, Patrick Bewick, Qi Li, Bingyu Zhao, Bo Zhang

## Abstract

The digestibility of soybean meal can be severely impacted by trypsin inhibitor (TI), one of the most abundant anti-nutritional factors present in soybean seeds. TI can restrain the function of trypsin, a critical enzyme that breaks down proteins in the digestive tract. Soybean accessions with low TI content have been identified. However, it is challenging to breed the low TI trait into elite cultivars due to a lack of molecular markers associated with low TI traits. We identified Kunitz trypsin inhibitor 1 (*KTI1*, Glyma01g095000) and *KTI3* (Glyma08g341500) as two seed-specific TI genes. Mutant *kti1* and *kti3* alleles carrying small deletions or insertions within the gene open reading frames were created in the soybean cultivar *Glycine max* cv. *Williams 82* (*WM82*) using the CRISPR/Cas9-mediated genome editing approach. The KTI content and TI activity both remarkably reduced in *kti1/3* mutants compared to the *WM82* seeds. There was no significant difference in terms of plant growth or maturity days of *kti1*/*3* transgenic and *WM82* plants in greenhouse condition. We further identified a T1 line, #5-26, that carried double homozygous *kti1/3* mutant alleles, but not the Cas9 transgene. Based on the sequences of *kti1*/*3* mutant alleles in #5-26, we developed markers to co-select for these mutant alleles by using a gel-electrophoresis-free method. The *kti1*/*3* mutant soybean line and associated selection markers will assist in accelerating the introduction of low TI trait into elite soybean cultivars in the future.

## Introduction

Soybean meal provides an excellent source of protein in animal feed since it is rich in amino acids with a high nutritional profile (Cromwell, Stahly et al. 1991). For instance, soy makes up 26% and 50% of swine and poultry feed, respectively (Gillman, Kim et al. 2015). However, reduced feed efficiency has been observed due to anti-nutritional and biologically active factors in raw soybean seeds (Liener 1996). Among these factors, trypsin inhibitor (TI) accounts for a substantial amount of this effect that cannot be ignored (Hymowitz 1986). TI restrains the activity of trypsin in monogastric animals. Because this enzyme is essential for optimal protein digestion, its restriction can lead to animal growth inhibition of 30-50% due to pancreatic hypertrophy/hyperplasia when raw soybeans are used in feed (Hymowitz 1986, Cook, Jensen et al. 1988, Liener 1994). In soybean meal processing facilities, TI in soybean meal is deactivated via a heating process at 90.5°C-100°C with the presence of 1% NaOH (Chen, Xu et al. 2014). This process not only reduces the nutritional value of soybean meal due to thermal destruction of amino acids, but also increases the energy cost of meal production by 25% (Chang, Tanksley et al. 1987).

With the recent increase in feed price and shipping cost, livestock farmers and grain operations are reconsidering soybean varieties with low-TI or TI-free traits as a way to reduce farm expenses by using raw soybeans as feed. Raising low-TI or TI-free soybeans on farms creates a niche market for integrated crop and livestock farmers, increasing their farm’s profitability. The use of soybean lines with genetically reduced levels of TI has proven as an effective strategy for improving animal growth. For instance, chicks fed soy-based diets with raw, unprocessed, low-KTI soymeal had higher feed efficiency ratios than chicks fed diets containing raw, unprocessed, conventional soybean meal (Batal and Parsons 2004). Thus, soybean cultivars with low TI content in the seeds is a long-term breeding goal for higher protein digestibility, better economic benefits, reduced environmental pollution caused by phosphorus, and the pursuit of sustainability for humanity and nature.

Plants have evolved a group of TI genes encoding proteins that can suppress the enzyme activities of proteases found in in plants, herbivores, animals and human beings (Jofuku and Goldberg 1989, Schuler, Poppy et al. 1999). The TIs in soybean can be classified into two families: the 21 kDa Kunitz trypsin inhibitor protein family (KTI) and the 7-8 kDa Bowman-Birk inhibitor protein family (BBTI) (Kunitz 1945, Wei 1983, Gillman, Kim et al. 2015). KTI proteins are thought to be largely specific for trypsin inhibition, while the major isoform of BBTI contains domains that interact with and inhibit both trypsin and chymotrypsin (Hwang, Foard et al. 1977, Gillman, Kim et al. 2015). Currently, only the KTI genes are targeted for selection of low-TI soybeans because KTI serves as the major contributor to trypsin inhibitor activity in soybeans. By far, the most significant success in reducing TI activity in soybean was the identification of a soybean accession (PI 157740) with dramatically reduced (∼40%) TI activity (Gillman, Kim et al. 2015). A frameshift mutation in *KTI3* (Gm08g341500) gene was identified in PI 157740, which is responsible for the low TI phenotype (Jofuku, Schipper et al. 1989). PI 157740 has been used in feeding trials, and it was found that raw extruded protein meal with lower KTI3 protein is superior for animal weight gain when compared to raw soybean meal harboring functional KTI3 (Cook, Jensen et al. 1988, Perez-Maldonado, Mannion et al. 2003). However, weight gain for young animals fed with non-heat-treated soybean materials including nonfunctional *KTI3* soybean materials is still inferior to those fed with heat-treated soybeans (Cook, Jensen et al. 1988, Perez-Maldonado, Mannion et al. 2003). Another soybean germplasm accession (PI 68679) was identified to carry a nonfunctional mutation on *KTI1* (Gm01g095000) gene (Gillman, Kim et al. 2015). *KTI1* and *KTI3* genes were determined to be synergistically controlling the TI content in soybean seeds (Gillman, Kim et al. 2015). Therefore, it is desirable to breed new soybean cultivars carrying both *kti1* and *kti3* mutant alleles. However, it is time-consuming to breed low TI soybean cultivars by selecting progenies derived from crosses between PI 157740, PI 68679, and elite varieties. Besides, the linkage drags associated with *KTI1* and *KTI3* may introduce undesirable agronomic traits, which could be difficult to remove by backcrossing. Although Kompetitive Allele Specific PCR (KASP) markers associated with *KTI3* and its mutant allele with 86% efficiency are available (Rosso, Shang et al. 2021), attempts at developing molecular markers associated with *KTI1* have not been successful (Gillman, Kim et al. 2015). Therefore, it is highly desirable to develop new *KTI1* mutant alleles that can be tagged with convenient molecular markers.

CRISPR/Cas9 mediated genome editing employs a Cas9 endonuclease and an 18-22 bp small guide RNA (sgRNA) that have a region that is complementary to a target gene sequence. The sgRNA binds to Cas9 and recruits the complex to target a gene. The Cas9 endonuclease generates DNA breaks, leading to mis-repaired target genes that contain deletions or insertions that disrupt gene function. In addition, several sgRNAs can be co-expressed in a single cell with Cas9, which allows the multiplex mutations of different genes simultaneously (Liang, Zhang et al. 2016). Because genome edited plants without transgenes are not considered as genetically modified organisms (GMO) (Kim and Kim 2016), mutant plants can be either directly released for field test or served as valuable resources for further breeding selection. Thus far, the CRISPR/Cas9 mediated genome editing technology has been widely used for targeted gene mutagenesis in diverse crop plant species, including soybean (Haun, Coffman et al. 2014, Jacobs, LaFayette et al. 2015, Cai, Chen et al. 2018), rice (Xu, Li et al. 2015), wheat (Upadhyay, Kumar et al. 2013), maize (Chen, Xu et al. 2018), tomato (Vu, Sivankalyani et al. 2020), cotton (Gao, Long et al. 2017), citrus (Peng, Chen et al. 2017), apple (Osakabe, Liang et al. 2018), grape (Osakabe, Liang et al. 2018), potato (Nakayasu, Akiyama et al. 2018), and banana (Shao, Wu et al. 2020) to improve their agronomic performances. For example, Jacobs et al. (2015) reported the first targeted mutagenesis in soybean using the CRISPR/Cas9 technology (Jacobs, LaFayette et al. 2015). Haun et al. (2014) generated a high oleic acid content soybean variety without transgenic components and improved the quality of soybean (Haun, Coffman et al. 2014). A soybean mutant with a late flowering phenotype was created using CRISPR/Cas9 technology to knock out the *GmFT2a* gene (Cai, Chen et al. 2018). Thus far, there is no research reporting an attempt to apply CRISPR/Cas9 to target anti-nutritional factors in soybean.

In this study, we aimed to (1) simultaneously knockout *KTI1* and *KTI3* genes in soybean cultivar Williams 82 via CRISPR/Cas9-mediated genome editing and (2) develop molecular markers associated with *kti1* and *kti3* mutant alleles that can be used for marker-assisted selection (MAS). We successfully recovered transgenic soybean plants that are carrying both *kti1* and *kti3* mutations. KTI content and trypsin inhibition activities (TIA) are dramatically decreased in the *kti1* and *kti3* mutant lines. In addition, we also developed molecular markers for co-selection of the new *kti1* and *kti3* mutant alleles. These *kti1* and *kti3* mutant lines and the newly developed selection markers have great potential for breeding the low TI trait into elite soybean varieties in the future.

## Results

### KTI gene family consists of multiple members with distinct expression patterns

Soybean *KTI* genes belong to a gene family with thirty-eight members (Soybase version Wm82.a4.v1) (Figure 1). As displayed in the genetic map, 38 *KTI* genes are located on 9 chromosomes (Chr) in the soybean genome. Specifically, Chr1 carries 2 *KTI* genes; Chr3 carries 1 *KTI* gene; Chr6 carries 1 *KTI* gene; Chr8 carries 13 *KTI* genes; Chr8 carries 10 *KTI* genes; Chr12 carries 1 *KTI* gene; Chr16 carries 8 *KTI* genes; Chr18 carries 1 *KTI* gene; and Chr19 carries 1 *KTI* gene (Figure 1).

**Figure 1.**
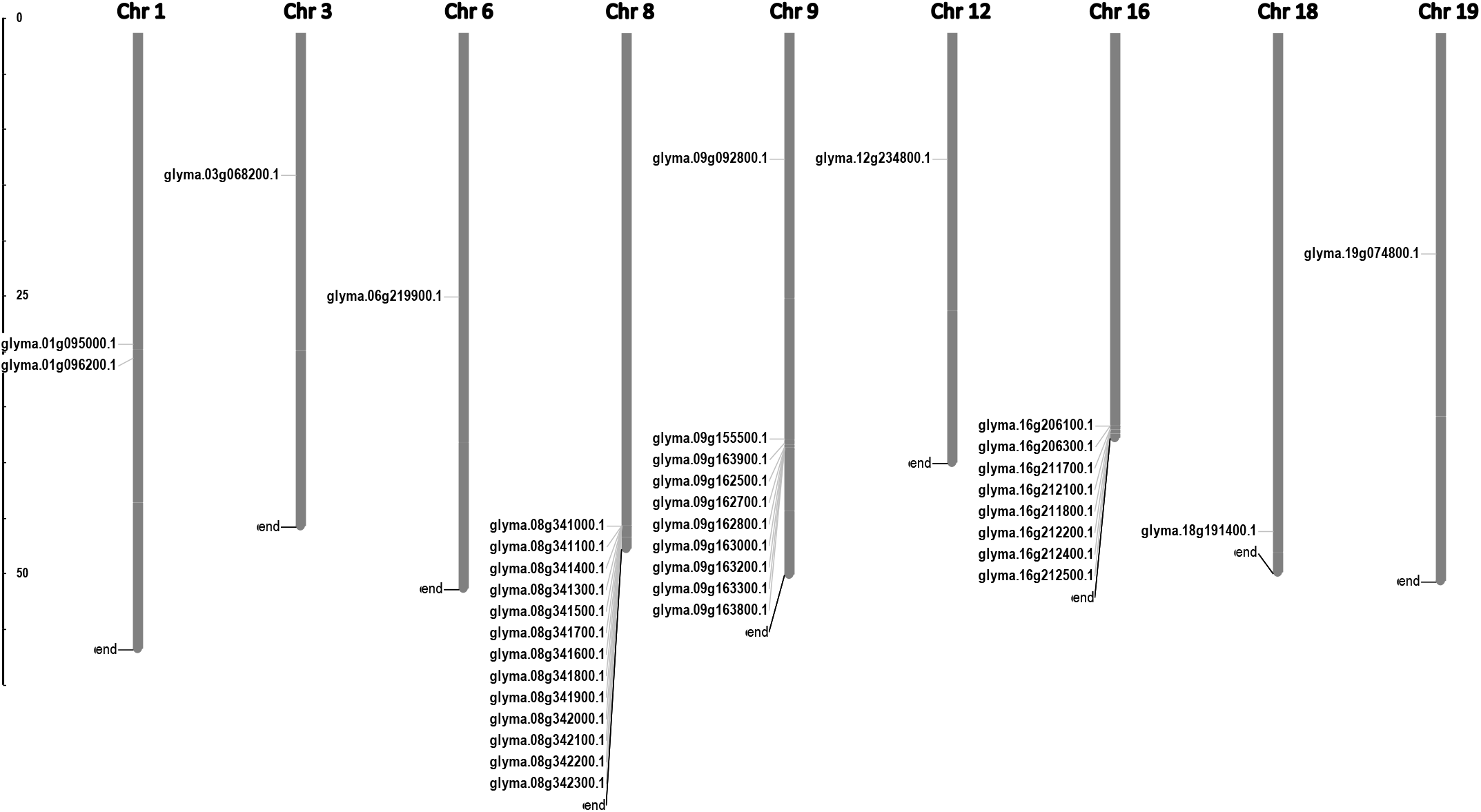
Physical mapping of 38 *GmKTI* genes on soybean 20 chromosomes. The gene map showing locations of *KTI* genes on soybean chromosomes was made using MapInspect. As displayed in the map, 38 *KTI* genes are located on 9 out of 20 chromosomes.

To identify the *KTI* genes expressed in the seed, we analyzed the expression patterns of all *KTI* genes in cv. *WM82* based on the expression data deposited in USDA ARS’s Soybase (Figure 2A). According to the expression patterns of *KTI* genes in various soybean tissues as displayed in Figure 2A, three *KTI* genes Gm01g095000 (*KTI1*), Gm08g341000, Gm08g342300, and Gm08g341500 (*KTI3*) were identified as seed-specific *KTI* genes. Soybean breeding lines V98-9005 (normal TI) and V03-5903 (low TI), presenting significantly different amounts of KTI concentration in seeds, were used to validate the tissue specific expressions of the four *KTI* genes by real-time PCR. Gm01g095000 (*KTI1*) and Gm08g341500 (*KTI3*) were predominately expressed in seeds compared to other tissues (Figure 2B). Both Gm08g341000 and Gm08g342300 had a relatively lower expression level in seeds than *KTI1* and *KTI3* but had higher expression in other tissue types (Figure 2B). Interestingly, both *KTI1* and *KTI3* had a relatively low expression level in the seeds of V03-5903 (low TI line), but higher expression in V98-9005 (normal TI line). Thus, we conclude that *KTI1* and *KTI3* are two major genes that may directly contribute to the TI contents in soybean seeds.

**Figure 2.**
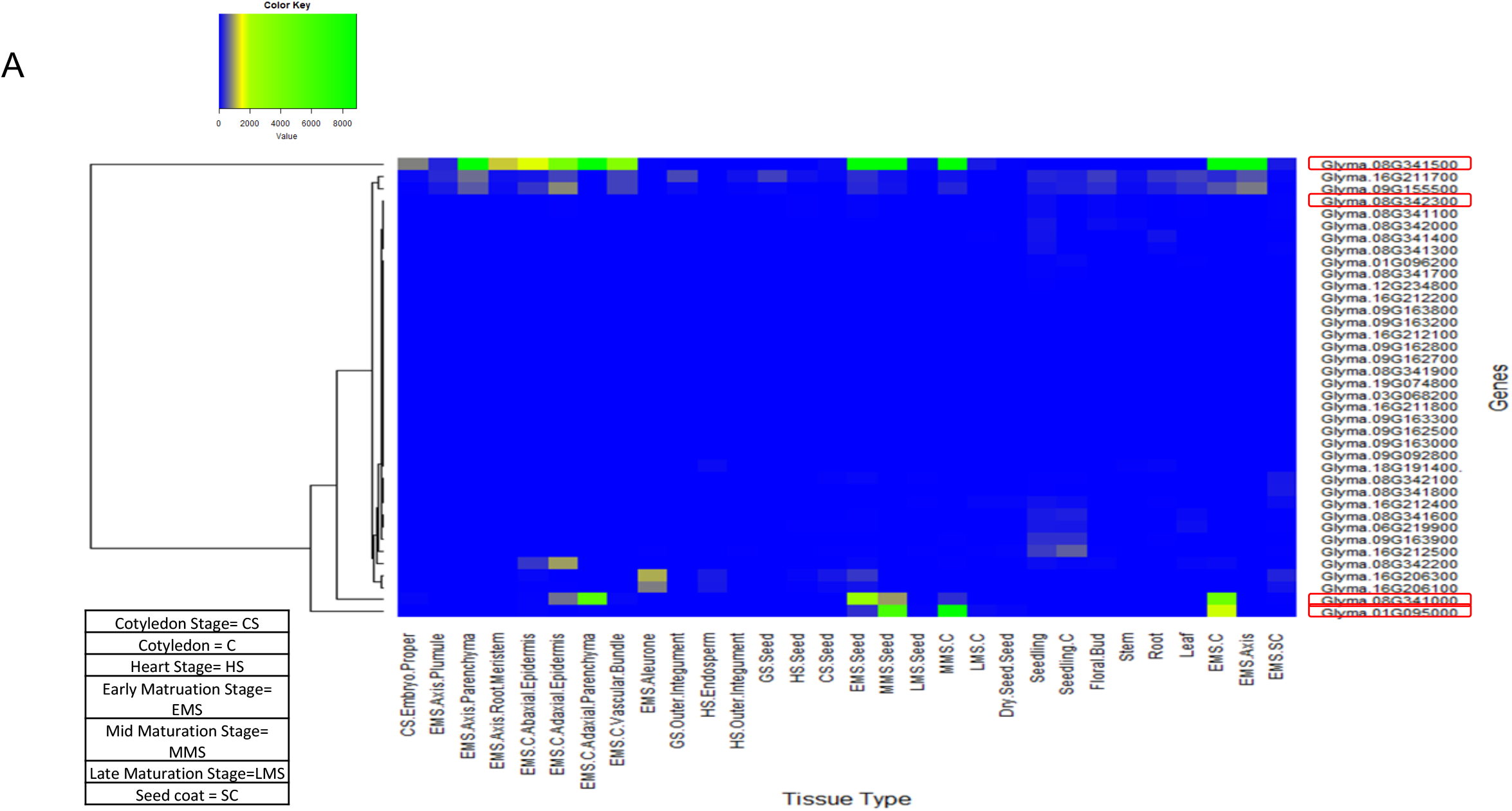
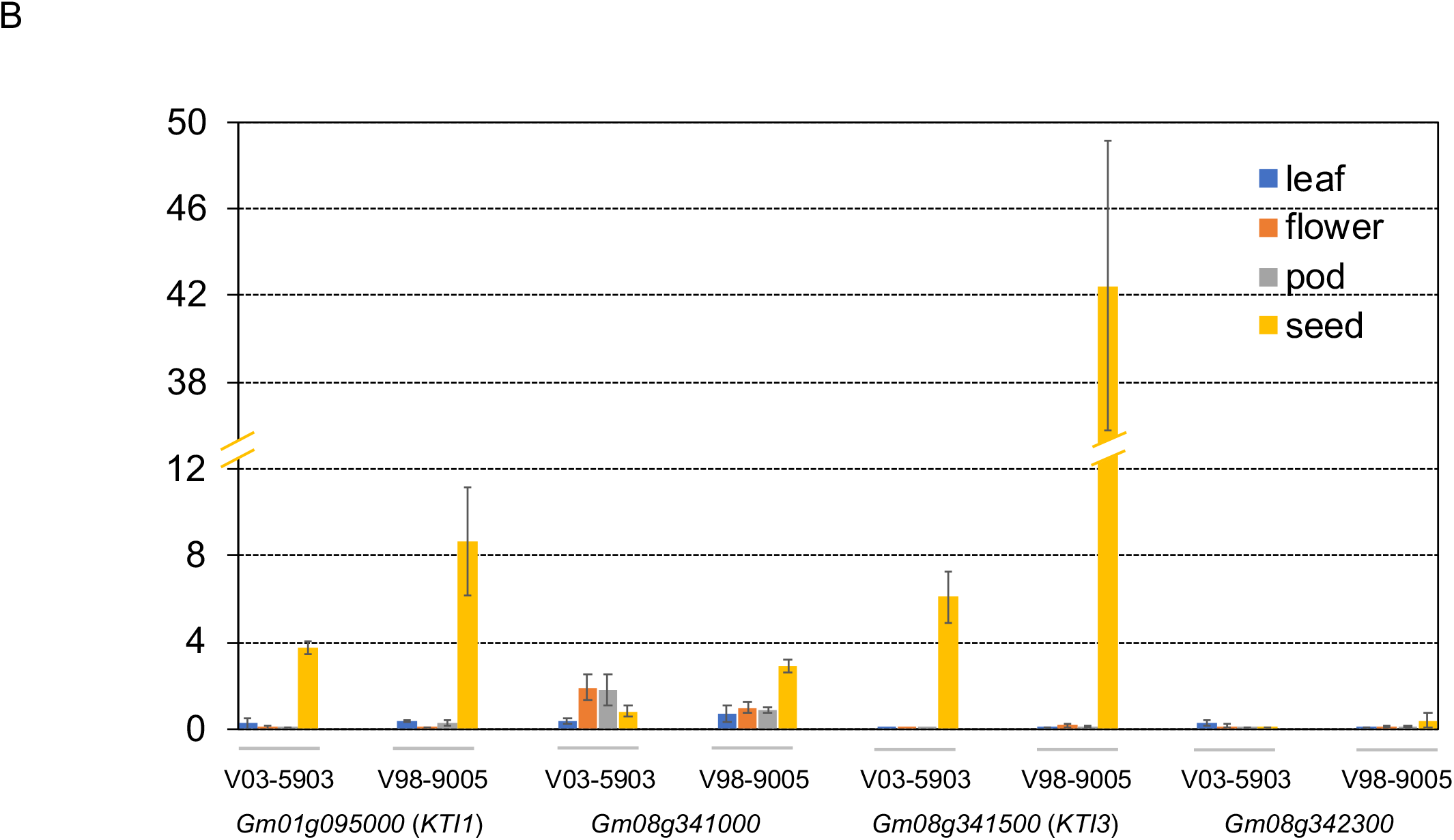
Expression levels of *KTI* genes in *WM82*. (A) RNA sequencing data of 38 *KTI* genes in 26 different tissue types of cv. *Williams 82* acquired from Phytozome soybean database was used to construct the heatmap to visualize their expression patterns. (B) The expressions of four soybean *KTI* genes were monitored by real-time PCR. Samples of leaf, flower, pod, and seed tissues from 2 breeding lines, V98-9005 (normal-TI line) and V03-5903 (low-TI line), were collected for RNA extraction. After reverse transcription, real-time PCR was used to evaluate the expressions of 4 genes including *Gm01g095000, Gm08g341000, Gm08g342300*, and *Gm08g341500* in different tissues with the ELF1B as the reference gene. The expression data was normalized as ΔCT and shown as mean ± s.e. Experiments were repeated three times and obtained similar results.

### Development of CRISPR/Cas9-based binary vector for genome-editing in soybean

To knock out the *KTI1* and *KTI3* genes from cv. *WM82* genome and create a new soybean cultivar with low TI content in soybean seeds, we developed a CRISPR/Cas9 construct, pBAR-Cas9-*kti13*, where the nuclease gene *Cas9* is expressed by Arabidopsis ubiquitin 10 (U10) promoter. A *bar* gene driven by a MAS promoter was used for selection of the putative transformants with Bialaphos or phosphinothricin (Figure 3A). A tandem array of two sgRNAs targeting *KTI1* and one sgRNA targeting *KTI3* was expressed by the U6 RNA promoter (Figure 3B).

**Figure 3.**
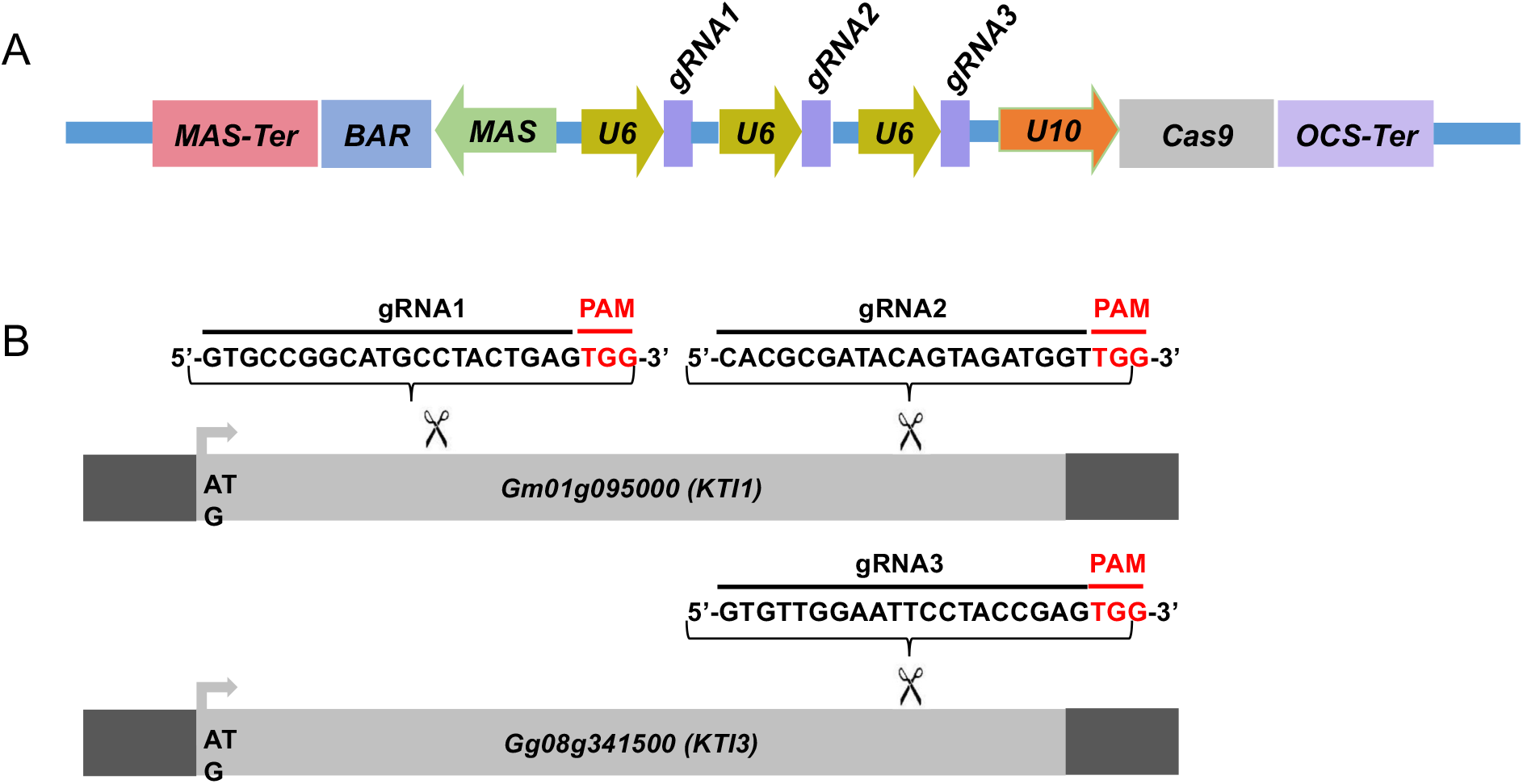
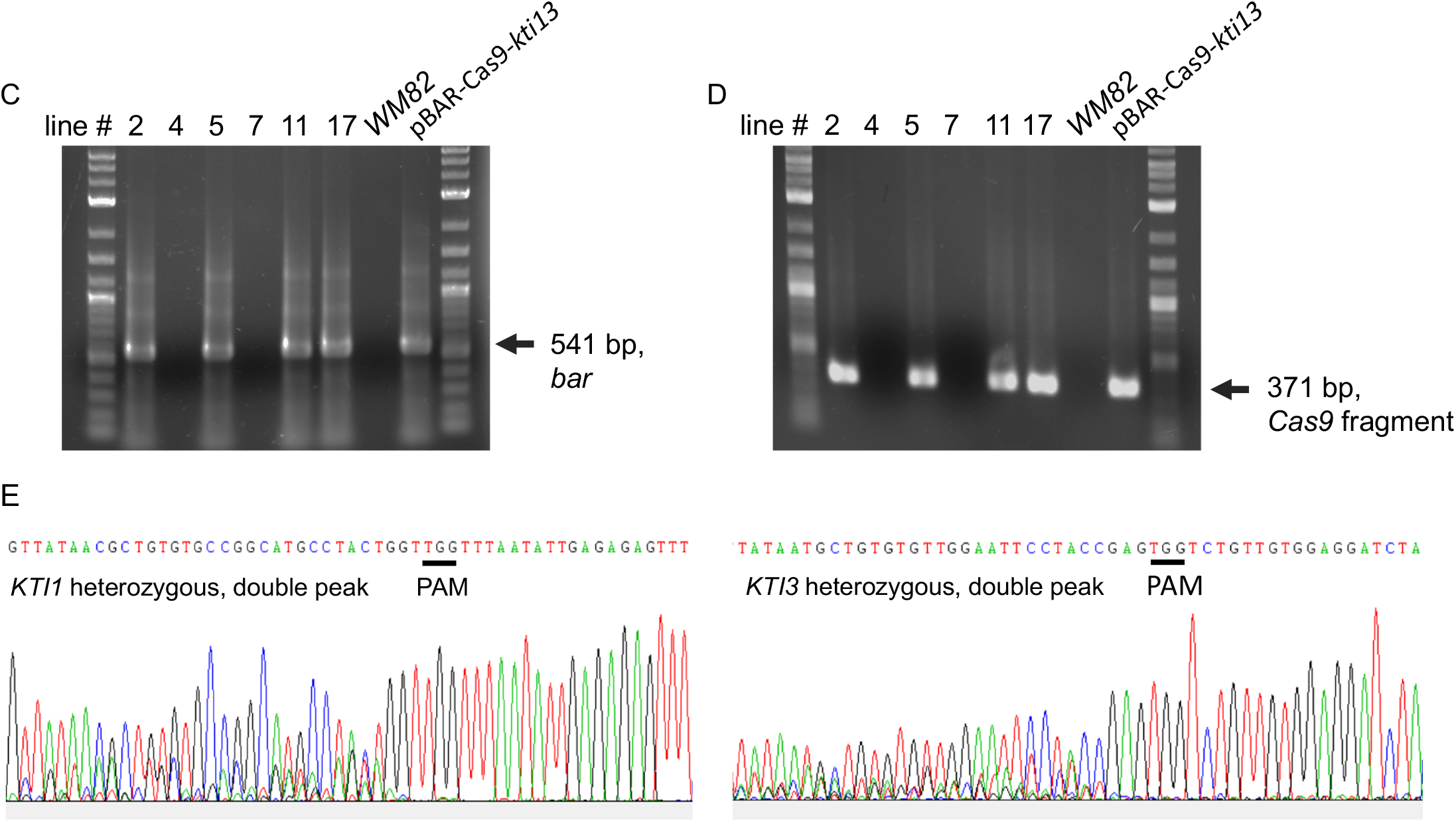
The scheme of binary vector used for CRISPR/Cas9 mediated gene editing on *KTI1*/*KTI3*, and transgenes and gene editing have been detected in the leaves of four T0 soybean plants. (A) The CRISPR/Cas9 construct harbors three necessary elements exhibited as below: the selection cassette consists of MAS promoter, *Bar* gene (soybean transformation selection marker), and MAS terminator; the Cas9 cassette consists of U10 promoter, Cas9 gene, and OCS terminator; three guide RNA cassettes and each of them consists of a U6 promoter, and one sgRNA. (B) The sequences of three sgRNAs is shown here. Two sgRNAs were designed, synthesized, and assembled to the plasmid to target on *KTI1*, while one sgRNA was designed, synthesized, assembled to the plasmid to target on *KTI3*. The fragments of two transgenes, (C) *Cas9* and (D) *Bar*, have both been detected in lines #2, #5, #11, and #17 by PCR, but not lines #4 and #7. The *WM82* gDNA serves as the template for negative control, while the plasmid DNA serves as the template for positive control. (E) The gene editing on *KTI1* and *KTI3* has also been observed in the leaf tissues of plants at T0 generation. The double peak sequence around the sgRNA region indicates the gene editing was ongoing but not completed.

### KTI1 and KTI3 genes are knocked out by CRISPR/Cas9 mediated gene editing

pBAR-Cas9-*kti1*/*kti3* was transformed into *WM82* via *Agrobacterium*-mediated transformation (Plant Transformation Facility at Iowa State University). Seventeen putative transgenic shoots were regenerated. Six shoots elongated and were transferred to rooting mediums. After further selection, they were transplanted into soil. Four lines, No. #2, #5, #11 and #17 were confirmed to be true transformants by positive amplification of the *bar* gene and a part of the *Cas9* gene (Figure 3C and D). The gene editing events in T0 plants were identified by amplification and sequencing of DNA fragments covering the sgRNA binding sites of *KTI1* and *KTI3*. The double peaks in the sequencing chromatograms suggest that both *KTI1* and *KTI3* genes were mutated and resulted in heterozygous alleles in the edited plant cells (Figure 3E). T0 seeds were harvested from T0 lines #2, #5, #11 and #17. Four T0 seeds of each line were randomly picked for DNA extraction and genotyping of the *KTI1* and *KTI3* genes via PCR amplification and DNA sequencing. The *KTI1* gene editing was completed and resulted in homozygous mutant alleles in all tested T0 seeds of the four lines. In addition, an identical gene editing pattern in *KTI1* was detected in all tested T0 seeds, in which a small DNA fragment (66bp) between two sgRNAs was lost after the gene editing (Figure 4A, B and C). Homozygous *KTI3* mutant alleles were only detected in T0 seeds from 2-3, #5-4, #11-2, and #11-4 (Figure 4D, E, F and G). The gene editing patterns in *KTI3* included both small deletions and insertions that all resulted in frameshift mutations in KTI3.

**Figure 4.**
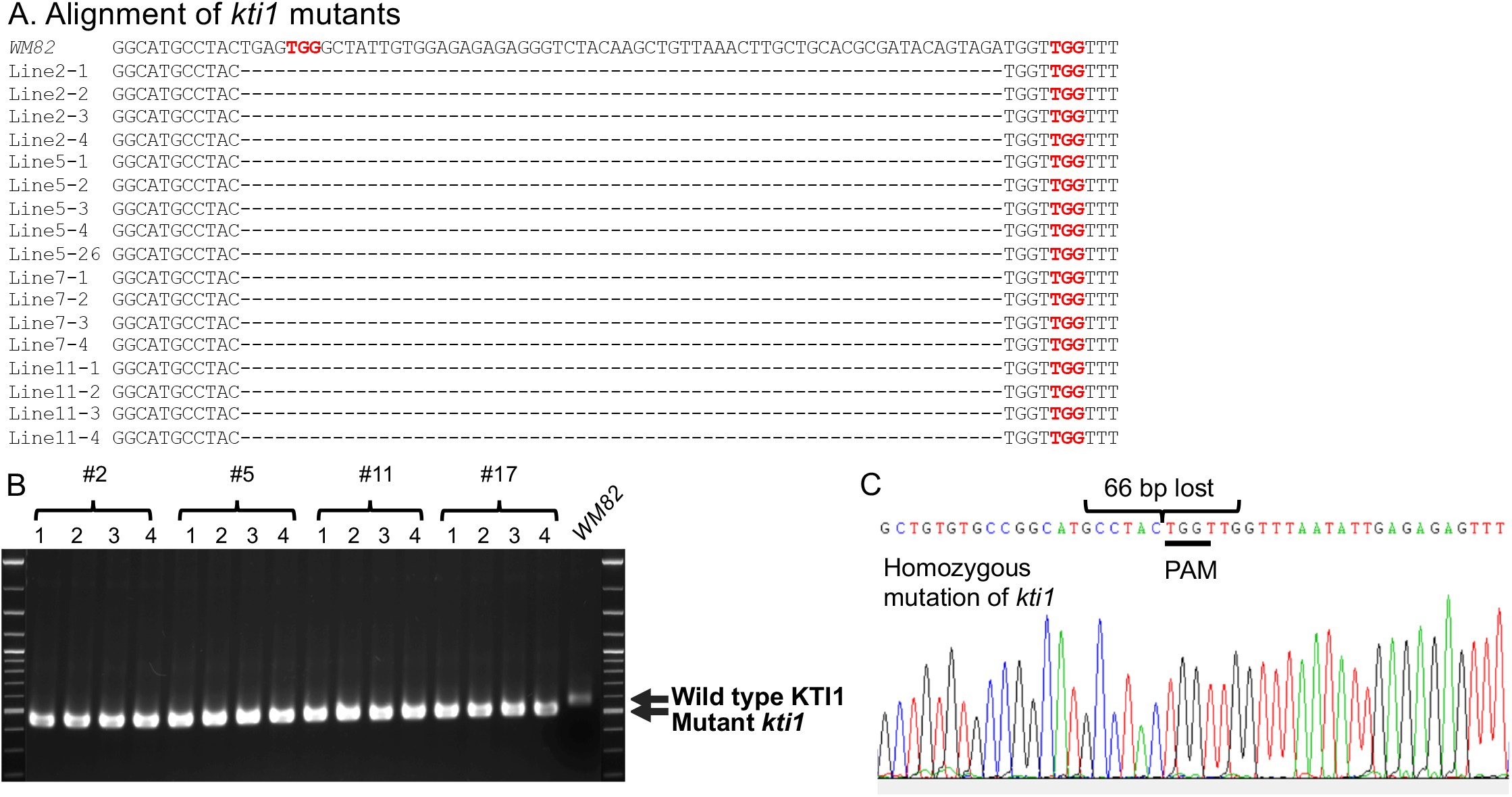
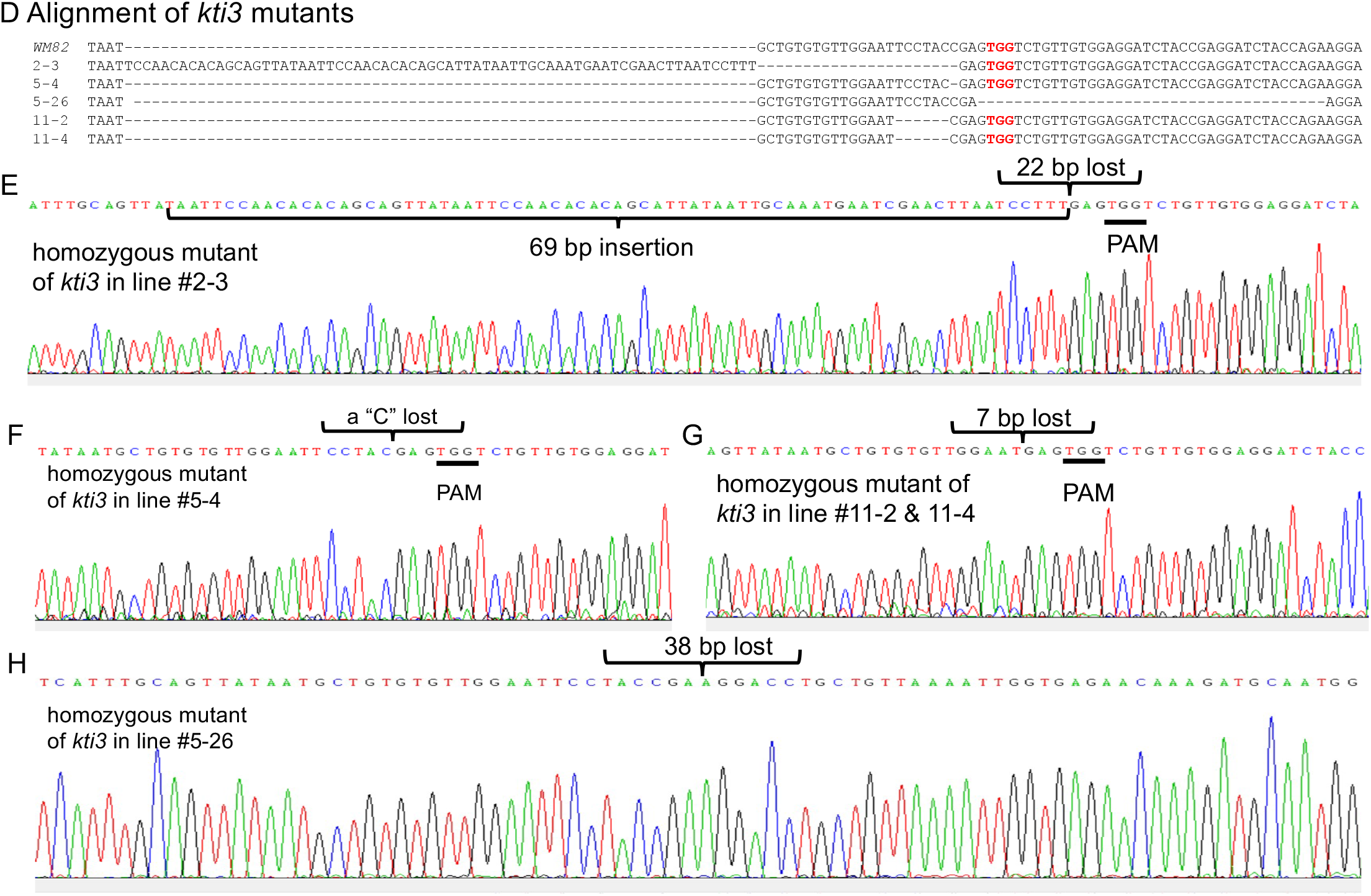
Gene editing on *KTI1* has been completed for all seeds of T0 generation while it has been completed on *KTI3* for some seeds of T0 generation. From each transgenic line (#2, #5, #11 and #17), four seeds of T0 generation were selected randomly for genotyping. (A) The alignment of mutant *kti1* in T0 seeds and T1 plant (#5-26) leaf, where the wild type *KTI1* in WM82 was the control. (B) Gel electrophoresis of *kti1* PCR products showed 16 seeds from line #2, #5, #11, and #17 had the same mutant on *kti1*, in which 66 nucleotides are lost between two sgRNAs. (C) Sanger sequencing result displayed the identical mutant *kti1*. (D) The alignment of mutant *kti3* in T0 seeds (#2-3, #5-4, #5-26, #11-2 and #11-4) and T1 plant (#5-26) leaf, where the wild type *KTI1* in WM82 was the control. (E), (F), (G), (H) showed the sanger sequencing results of *kti3* mutant in #2-3, #5-4, #11-2, #11-4, and #5-26.

### TI content and activity dramatically declined in the edited soybean seeds

T0 seeds were also used for quantification of the KTI content by using a HPLC-based approach (Rosso, Shang et al. 2018). The tested seeds of #2-3, #5-4, #11-2, and #11-4, which carried mutations on both *KTI1* and *KTI3* genes had the lowest KTI content (Figure 5). The tested seeds of #2-1, #5-1, #11-1, and #17-1, with only the *KTI1* mutation, also had lower KTI content than the wild-type *WM82* seeds (Figure 5). The KTI content in other genotyped seeds with editing only on *KTI1* was also lower than that in *WM82* seeds (data not shown). We further tested the trypsin inhibition activity (TIA) using crude protein extracts from the T0 seeds. As shown in Figure 6, the crude proteins of seeds with mutant *kti1* and *kti3* had the lowest TIA (Figure 6). The seeds with mutant *kti1* only also had reduced TIA (Figure 6) in comparison with *WM82* and Glenn (a commercial soybean cultivar as a control). The KTI content and TIA were ranked in order as: *kti1*/*3* double mutant < *kti1* single mutant ≤ PI 547656 (low TI accession) < *WM82* < Glenn. Taken together, we conclude that *KTI1* and *KTI3* are two major genes responsible for the KTI content and TIA in soybean seeds. Therefore, knockout of *KTI1* and *KTI3* reduced the KTI content and impaired the TIA in soybean seeds.

**Figure 5.**
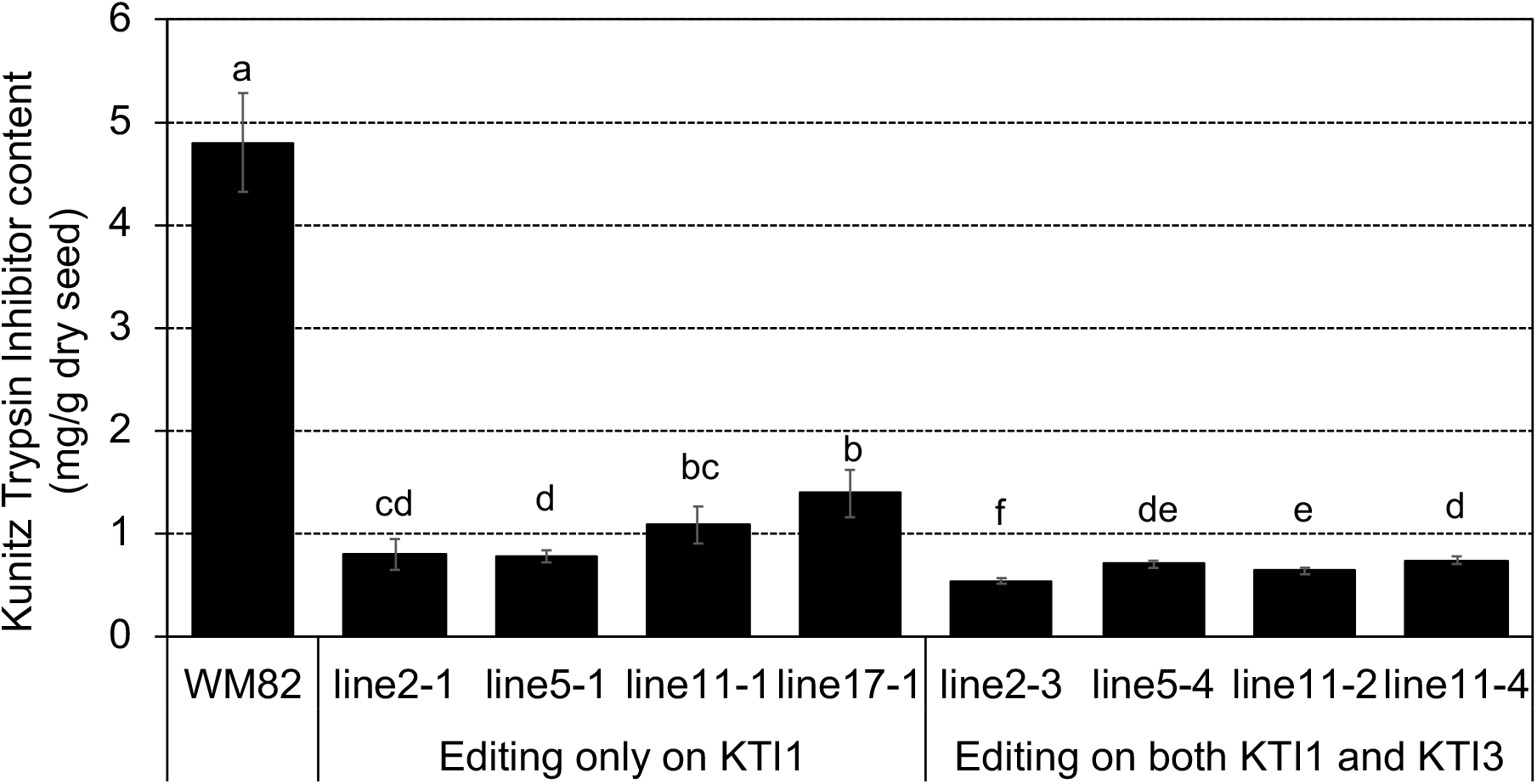
KTI content declined dramatically in gene-edited seeds. KTI content was measured in 4 double mutated seeds (#2-3, #5-4, #11-2, and #11-4) and 4 seeds with a single mutation on *KTI1* (#2-1, #5-1, #11-1, and #17-1), where the KTI content in *WM82* seed served as the control. Experiments were conducted with three technical replicates and showed comparable results, shown as mean ± s.e. Different letters indicate significant differences.

**Figure 6.**
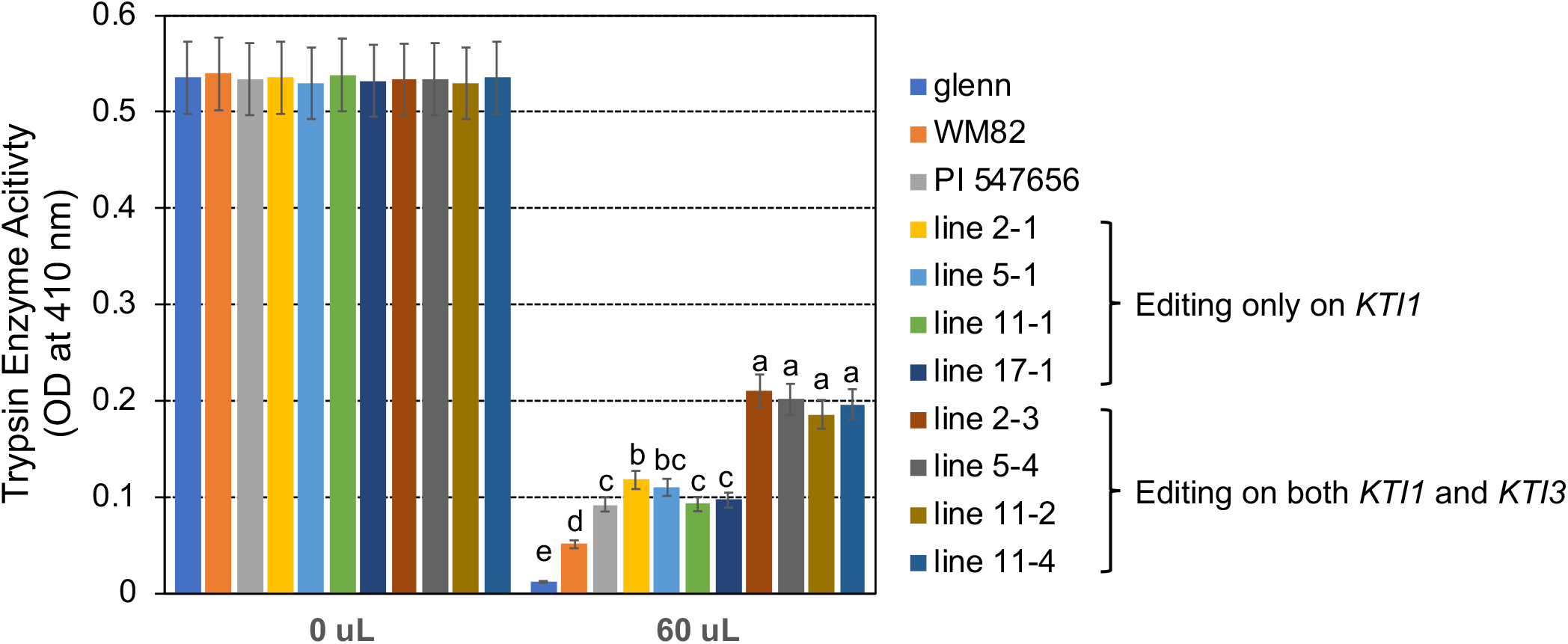
TIA declined dramatically in the gene-edited seeds. Bovine trypsin enzyme activities were measured using crude extracts of 4 double mutated seeds (#2-3, #5-4, #11-2, and #11-4), 4 seeds with a single mutation on *KTI1* (#2-1, #5-1, #11-1, and #17-1), and *WM82*. Experiments were conducted with three technical replicates and showed comparable results, shown as mean ± s.e. Different letters indicate significant differences.

The edited *KTI1* gene lost 66 bp that may result in mutant proteins with deletion of 22 amino acids. Truncated *KTI1* may still possess some TIA. To rule out this possibility, we also tested the TIA of truncated KTI1_Δ22aa_ protein *in vitro*. To this end, we cloned the open reading frames of *KTI1*_Δ66bp_ and wild-type *KTI1* and *KTI3* into a protein expression vector, in which a 6xHis tag is fused to C-terminus of the expressed proteins. The purified proteins were subjected to a TIA assay which showed that while KTI1 and KTI3 both could inhibit trypsin activity, the truncated KTI1_Δ22aa_ failed to suppress trypsin activity (Figure S1A and B). Therefore, the new *kti1* allele (*KTI1*_Δ66bp_) encodes a truncated protein that loses its TI function.

### Knockout KTI1 and KTI3 did not affect plant growth and maturity period days of soybean

To examine whether the mutant *kti1/3* could significantly affect plant growth and maturity period days of soybean, we planted T0 seeds of line #2 and #5, and *WM82* in a greenhouse. By Bialaphos-mediated screening, we classified the T1 plants from line #2 and #5 as transgene-free plants or transgenic plants. We measured the agronomic traits of the transgenic plants including plant height, the number of main branches per plant, number of pods bearing branches, number of pods, leaf length, leaf width and petiole length. There was no significant difference in terms of all measured agronomic traits among the plants of *WM82*, Line 2 and Line 5 (Table 1). We also measured the maturity period days of the soybean plants by recording the dates from planting to beginning bloom (R1), to beginning pod (R3), to beginning seed (R5), to full seed (R6), to maturity (R8), and the total lifespan (from planting to maturity). There were no remarkable differences in terms of R1, R3, R5, R6, R8 and total life span among all tested plants (Table 1). Therefore, we conclude that knockout of *KTI1* and *KTI3* did not alter plant growth or the maturity period of soybean lines tested.

**Table 1.**
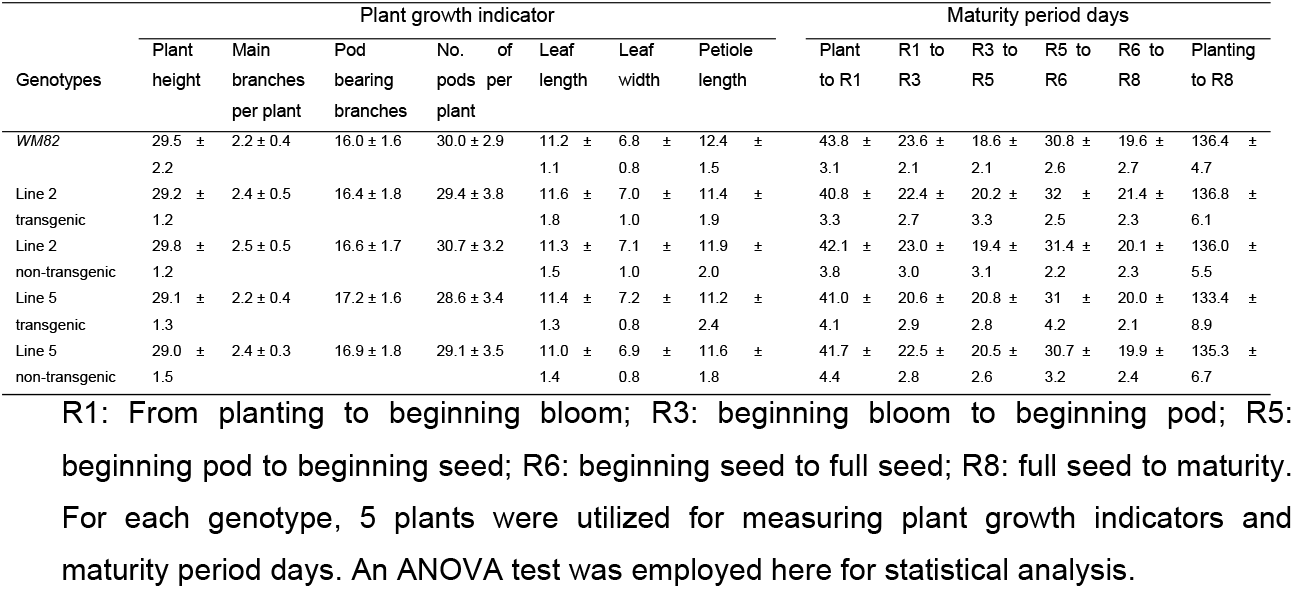
Knockout of *KTI1* and *KTI3* does not alter the plant growth indicator and maturity period days of cv. *WM82*.

| Genotypes | Plant growth indicator |  |  |  |  |  |  |  | Maturity period days |  |  |  |  |  |
| --- | --- | --- | --- | --- | --- | --- | --- | --- | --- | --- | --- | --- | --- | --- |
|  | Plant height | Main branches per plant | Pod bearing branches | No. of pods per plant | Leaf length | Leaf width | Petiole length |  | Plant to R1 | R1 to R3 | R3 to R5 | R5 to R6 | R6 to R8 | Planting to R8 |
| *WM82* | 29.5 ± 2.2 | 2.2 ± 0.4 | 16.0 ± 1.6 | 30.0 ± 2.9 | 11.2 ± 1.1 | 6.8 ± 0.8 | 12.4 ± 1.5 |  | 43.8 ± 3.1 | 23.6 ± 2.1 | 18.6 ± 2.1 | 30.8 ± 2.6 | 19.6 ± 2.7 | 136.4 ± 4.7 |
| Line 2 transgenic | 29.2 ± 1.2 | 2.4 ± 0.5 | 16.4 ± 1.8 | 29.4 ± 3.8 | 11.6 ± 1.8 | 7.0 ± 1.0 | 11.4 ± 1.9 |  | 40.8 ± 3.3 | 22.4 ± 2.7 | 20.2 ± 3.3 | 32 ± 2.5 | 21.4 ± 2.3 | 136.8 ± 6.1 |
| Line 2 non-transgenic | 29.8 ± 1.2 | 2.5 ± 0.5 | 16.6 ± 1.7 | 30.7 ± 3.2 | 11.3 ± 1.5 | 7.1 ± 1.0 | 11.9 ± 2.0 |  | 42.1 ± 3.8 | 23.0 ± 3.0 | 19.4 ± 3.1 | 31.4 ± 2.2 | 20.1 ± 2.3 | 136.0 ± 5.5 |
| Line 5 transgenic | 29.1 ± 1.3 | 2.2 ± 0.4 | 17.2 ± 1.6 | 28.6 ± 3.4 | 11.4 ± 1.3 | 7.2 ± 0.8 | 11.2 ± 2.4 |  | 41.0 ± 4.1 | 20.6 ± 2.9 | 20.8 ± 2.8 | 31 ± 4.2 | 20.0 ± 2.1 | 133.4 ± 8.9 |
| Line 5 non-transgenic | 29.0 ± 1.5 | 2.4 ± 0.3 | 16.9 ± 1.8 | 29.1 ± 3.5 | 11.0 ± 1.4 | 6.9 ± 0.8 | 11.6 ± 1.8 |  | 41.7 ± 4.4 | 22.5 ± 2.8 | 20.5 ± 2.6 | 30.7 ± 3.2 | 19.9 ± 2.4 | 135.3 ± 6.7 |

R1: From planting to beginning bloom; R3: beginning bloom to beginning pod; R5: beginning pod to beginning seed; R6: beginning seed to full seed; R8: full seed to maturity. For each genotype, 5 plants were utilized for measuring plant growth indicators and maturity period days. An ANOVA test was employed here for statistical analysis.

### Development of molecular markers for selection of the kti1 and kti3 alleles

A double homozygous *kti1* and *kti3* mutant plant #5-26 that did not carry the Cas9 transgene was selected from T1 generation plants (Figure 4A, C, D and H). The ‘transgene-free’ soybean plants can be used to breed the low TI trait into other elite soybean cultivars. In order to co-select the *kti1* and *kti3* mutant alleles in the derived progenies, we attempted to develop co-dominate molecular markers that can distinguish between wild-type *KTI1*/*KTI3* and mutant *kti1*/*kti3* alleles.

In the genome of #5-26, the mutant allele of *kti1* had a 66 bp deletion. We designed three PCR primers, ZW1, ZW2 and ZW3 (Figure 7A and Table S1). ZW1 was a common reverse primer that can bind to the same region in both *KTI1* and *kti1* alleles, while ZW2 and ZW3 were both forward primers binding to unique sequences of *KTI1* and *kti1* alleles, respectively (Figure 7A). Soybean cultivars that carry the wild-type *KTI1* gene (*WM82*) amplified a 180 bp DNA fragment when hybridized with ZW1 and ZW2, but failed to amplify any fragments when hybridized with ZW1 and ZW3. On the contrary, the soybean lines carrying a homozygous *kti1* mutant allele (#5-26) amplified a 134 bp DNA fragment with ZW1 and ZW3, but failed to amplify any fragments with ZW1 and ZW2 (Figure 7B).

**Figure 7.**
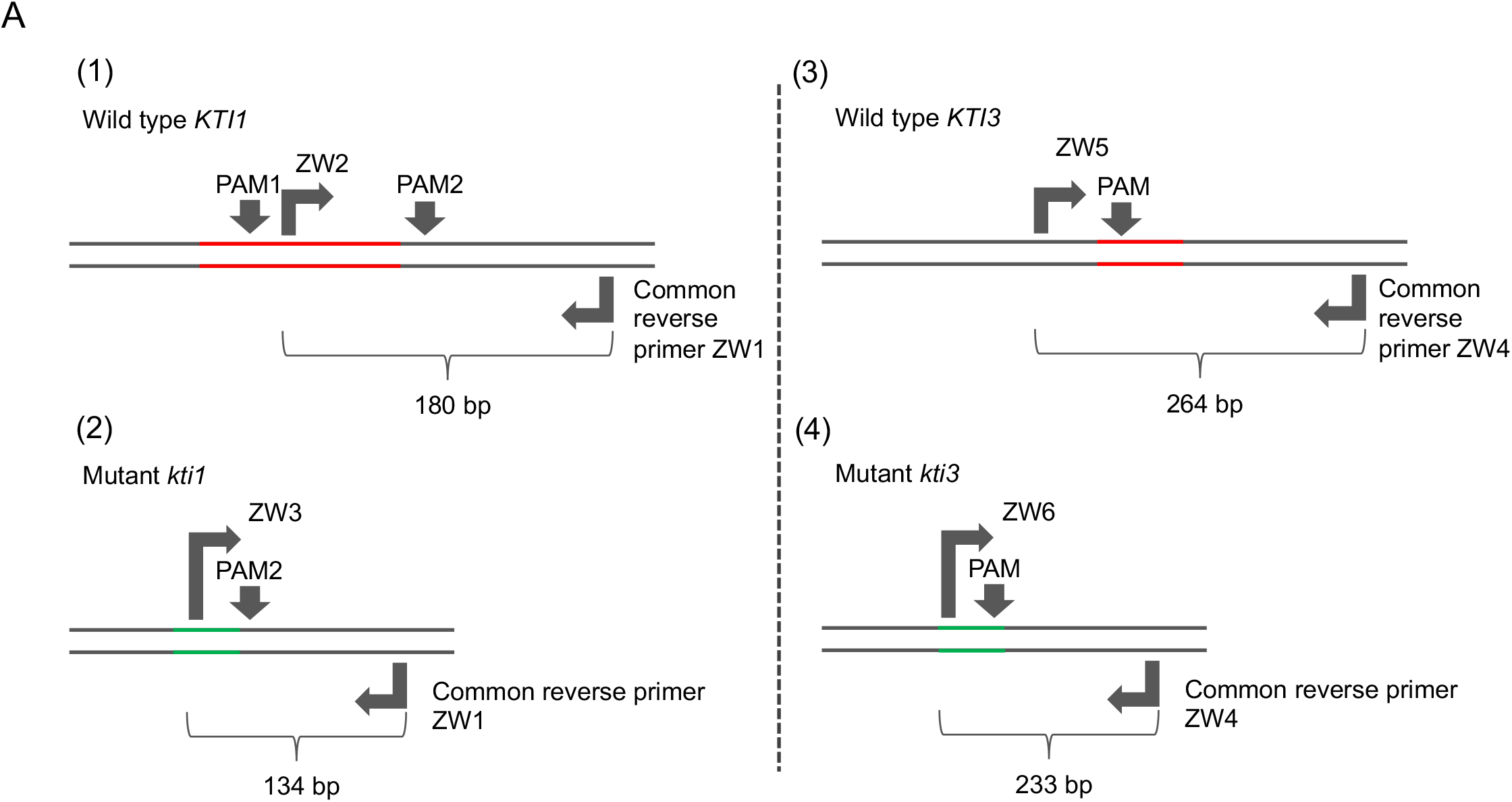
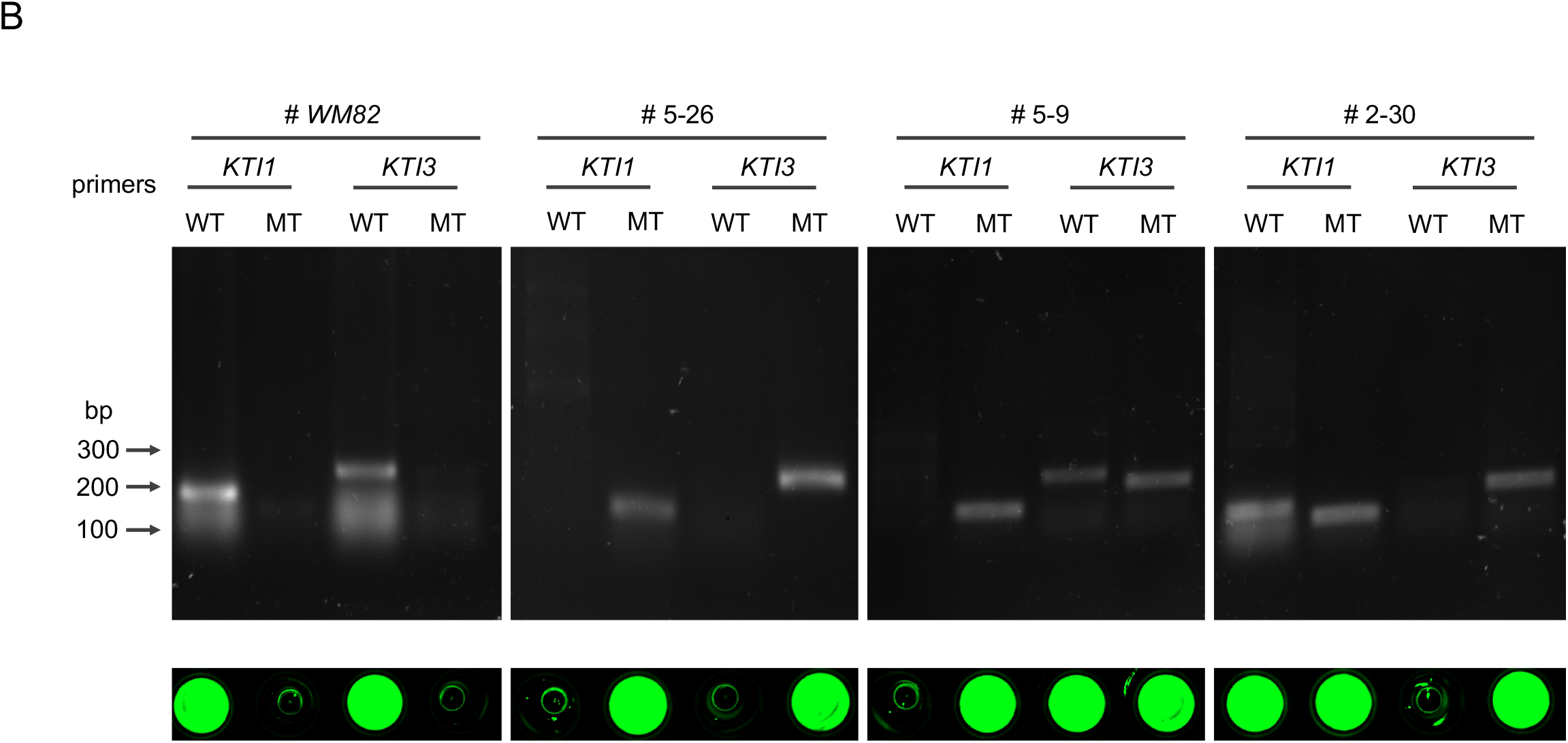
The development of selection markers for breeding low KTI soybean varieties based on the *kti1* and *kti3* mutants generated by CRISPR/Cas9-mediated gene editing. (A) Schematic development of primers for amplification of wild type *KTI1* (1), mutant *KTI1* (2), wild type *KTI3* (3), and mutant *KTI3* (4). The red lines indicate the lost fragment in *KTI1* or *KTI3* during gene editing. The green lines indicate new DNA regions in *kti1* or *kti3* generated by splicing two fragments. (B) The 4 pairs of primers in (A) were utilized to amplify the alleles of *KTI1, kti1, KTI3*, and *kti3* with gDNA of four different soybean genotypes, including *WM82*, three transgenic lines #5-26, #5-9, and #2-30. Based on our genotyping data, #5-26 has homozygous mutations of *kti1* and *kti3*; #5-9 only has a homozygous mutation of *kti1* but carries the heterozygous mutation of *kti3*; #2-30 only has a homozygous mutation of *kti3* but carries the heterozygous mutation of *kti1*. Thus, it was clear that the pair of ZW1/ZW2 can amplify wild type *KTI1* from *WM82* and #2-30 gDNA in PCR tests, while the pair of ZW1/ZW3 can amplify mutant *kti1* from #5-9 and #5-26 gDNA. Also, the pair of ZW4/ZW5 can amplify wild type *KTI3* from *WM82* and #5-9 gDNA, while ZW4/ZW6 can amplify mutant *kti3* from #2-30 and #5-26 gDNA. As shown in the bottom panel, only the positive PCR products incubated with the dye of sybrgreen at 75 °C can display the fluorescent signals, suggesting the reliability of the developed gel-electrophoresis-free method for screening mutant alleles of *kti1* and *kti3*.

In the genome of line #5-26, the mutant *kti3* allele had a 38 bp deletion, which allowed us to design PCR primers ZW4, ZW5 and ZW6 (Figure 7A and Table S1). ZW4 was a common reverse primer for both *KTI3* and *kti3*, while ZW5 and ZW6 were forward primers matched with unique sequences of *KTI3* and *kti3*, respectively (Figure 7A). A soybean cultivar carrying the wild-type *KTI3* gene (*MW82*) amplified a 264 bp DNA fragment with primers ZW4 and ZW5, but not with primers ZW4 and ZW6. In contrast, the soybean line carrying homozygous *kti3* allele (#5-26) can amplify a 233 bp DNA fragment with primers ZW4 and ZW6, but not ZW4 and ZW5 (Figure 7B).

We further tested these PCR primers by amplifying DNA fragments from two T1 plants that were genotyped by DNA sequencing. Line #5-9 had homozygous *kti1* alleles and heterozygous *KTI3/kti3* alleles, where the *kti3* allele was identical to the one in #5-26. Line #2-30 had homozygous *kti3* alleles and heterozygous *KTI1*/*kti1* alleles, where the *kti1* allele was identical to the one in #5-26. As shown in Figure 7B, PCR amplification with the different combinations of ZW1, ZW2, ZW3, ZW4, ZW5 and ZW6 can accurately identify the *KTI1/kti1* and *KTI3/kti3* genotypes of #5-9 and #2-30 (Figure 7B). Therefore, we successfully developed molecular markers to select the *kti1* and *kti3* mutant alleles generated by CRISPR/Cas9 mediated mutagenesis. These molecular markers can assist in the breeding selection of low TI soybean plants harboring *kti1*/*3*.

To simplify the procedure of marker-aided selection, we tested a gel-electrophoresis-free protocol that can be implemented for high throughput screening of progenies derived from a cross between a soybean cultivar carrying wild-type *KTI1*/*3* and one carrying the *kti1*/*3* mutant. In brief, all PCR products as described above were mixed with 1X SYBR Green and heated at 75°C for 10 mins and then visualized under UV light. As shown in Figure 7B, the fluorescent signals were the indications of positive amplifications in WM82 with primers ZW1/ZW2 and ZW4/ZW5, while in #5-26, the fluorescent signals can only be observed with primers ZW1/ZW3 and ZW4/ZW6 (Hirotsu, Murakami et al. 2010). Therefore, we identified the homozygous *kti1* and *kti3* alleles by directly staining the PCR products without the need of gel electrophoresis, which can significantly reduce the cost of labor and time.

## Discussion

In this study, we optimized a CRISPR/cas9-vector for genome editing in soybean (Figure 3A). The modified vector allowed us to simultaneously knock out two seed specific KTI genes (*KTI1* and *KTI3*). The *kti1*/*3* mutant plants grew normally in greenhouse conditions, and the seeds of *kti1*/*3* mutant had dramatically reduced KTI content and TI activities in comparison with wild type seed of *WM82*.

Soybean is one of the important sources of protein for animal and human consumption. However, in their evolution, soybeans have developed diverse defense components to protect seeds from being eaten by insects and animals including trypsin inhibitor, phytate acid, and raffinose family of oligosaccharides (RFOs). In the agricultural practice, the anti-nutritional and biologically active factors are responsible for reduced feed efficiency when raw soybeans are fed to animals. Therefore, it is of great significance to increase feed efficiency, especially the protein digestibility via assembling gene function exploration, application, and advanced genetically engineering together into the soybean industry. Proteinaceous plant trypsin inhibitors are a diverse family of (poly)peptides that play diverse roles in plant growth such as maintaining physiological homeostasis and serving the innate defense machinery (Li, Brader et al. 2008, Junker, Zeissig et al. 2012, Arnaiz, Talavera-Mateo et al. 2018, Zhao, Ullah et al. 2019). Since TI proteins exert direct effects on pests and herbivores by interfering with their physiology, any food containing TI proteins will be avoided by these organisms. In alfalfa, the trypsin inhibitors Msti-94 and Msti-16 were demonstrated to act as a stomach poison, significantly reducing the survival and reproduction rates of aphid (Zhao, Ullah et al. 2019). Mutant plants with reduced TI are usually more susceptible to pests. In wheat, α-amylase/trypsin inhibitors (ATIs) CM3 and 0.19 were identified as pest-resistance molecules, activating innate immune responses in monocytes, macrophages, and dendritic cells (Junker, Zeissig et al. 2012). For example, the *Arabidopsis* lines containing silenced *atkti4* and *atkti5* were found to have a higher susceptibility to *T. urticae* (Spider mite) than wild-type plants (Arnaiz, Talavera-Mateo et al. 2018). RNAi silencing of the *AtKTI01* gene resulted in enhanced lesion development after infiltration of leaf tissue with the programmed cell death eliciting fungal toxin fumonisin B1 or the avirulent bacterial pathogen *Pseudomonas syringae* pv. *tomato DC3000* carrying *avrB* (Li, Brader et al. 2008). Although similar defense functions have not been reported on TI genes in the soybean genome, it is reasonably suspected that certain members of the KTI gene family have comparable protecting roles for soybean plants.

As previously discussed, the high concentration of TI proteins in soybean meal restricts the function of trypsin, which causes low digestibility and reduces its nutritional value. Thus, cultivars have been developed by introgression of low TI traits into elite cultivars. We previously developed a low TI line via conventional breeding: V12-4590. During field trials in 2017 and 2018, we observed that this low TI line is indeed more susceptible to multiple phytopathogens such as: all races of Soybean Cyst Nematode (SCN) (*Heterodera glycines*), Stem Canker (*Diaporthe aspalathi*), Cercospora leaf blight (*Cercospora kukuchii*), Soybean vein necrosis virus, and Downy Mildew (*Peronospora manshurica*) (Zhang, unpublished data). This suggests that that the soybean *KTI* genes that are negatively selected do indeed have a role in plant immunity. It is also possible that some plant immunity related genes that genetically link with *KTI* genes are negatively selected during the breeding process. As shown in Figure 1, at least 13 KTI genes are clustered in a small region on chromosome 8. Interestingly, we also identified a putative TGACG-Binding (TGA) transcription factor (TF) that is tightly linked to the KTI gene cluster at chromosome 8. Arabidopsis TGA TFs play a positive role in systemic acquired resistance (SAR) that is crucial in plant immunity (Hussain, Sheikh et al. 2018). Therefore, breeding of low TI soybean lines resulting in the loss or mutation of both of the KTI genes and the TGA TF gene, leading to an increased susceptibility of soybean challenged by phytopathogens.

The *kti1*/*3* mutant soybean line generated via CRISPR/Cas9-mediated mutagenesis is an isogenic line of wild type *WM82* (table 1). Therefore, it will be an ideal test subject to see if *KTI1*/*3* has a direct role in plant immunity. Since *KTI1*/*3* were almost only expressed in seeds (Figure 2) (Gillman, Kim et al. 2015), the knockout of these two genes may not interfere with plant immunity in non-seed tissue, which deserves to be further investigated in the future.

In this study, we identified a *kti1*/*3* double homozygous mutant along with *kti1* and *kti3* single homozygous mutants. The *kti1*/*3* double homozygous mutant has the lowest KTI content and trypsin inhibition activity (Figure 5 & 6). Therefore, it can be determined that (Hussain, Sheikh et al. 2018)*KTI1* and *KTI3* synergistically contribute to the KTI content and TI activity in soy proteins. The previous report suggests that the soybean line carrying natural mutations of *kti1* and *kti3* has increased BBTI content (Gillman, Kim et al. 2015). It is unclear if the increased BBTI content is caused by un-intentional selection during the breeding process or if the expression of *BBTI* genes is increased because of the mutations of two *KTI* genes. Therefore, it will be interesting to test the BBTI content and activity in the seed proteins of the *kti1*/*3* mutant generated in this study.

Despite the fact that the CRISPR/Cas9 technique has been successfully utilized to generate various soybean mutants (Jacobs, LaFayette et al. 2015), the current *agrobacterium*-mediated soybean transformation protocol is inefficient and genotype dependent. This limits the wide implementation of CRISPR/Cas9 technique in soybean breeding programs (Yamada, Takagi et al. 2012). The soybean transformation protocol employs Bialaphos as the selection agent (Luth, Warnberg et al. 2015). The *Bar* gene is used as the selection marker gene and encodes a phosphinothricin acetyltransferase protein that can confer the transformants’ resistance to bialaphos. It has been reported that the *Bar* gene expression must be fine-tuned in order to successfully select true transgenic plants (Testroet, Lee et al. 2017). The original CRISPR/Cas9 vector, pCut, has used a MAS (mannopine synthase) promoter to express the *Bar* gene (Peterson, Haak et al. 2016). However, for unknown reasons, the vector does not work well, even in *Arabidopsis thaliana* (Liu and Zhao, unpublished data).

With the intention of improving the transformation system, we modified the Bialaphos selection vector in the pMU3T (Liu, Miao et al. 2016). Specifically, we replaced the Kanamycin selection marker gene with the *Bar* gene, whose expression was driven by a MAS promoter (Figure 3A). The MAS promoter is known to be most active in the roots of emerging seedlings and very active in the cotyledons and lower leaves (Langridge, Fitzgerald et al. 1989). Despite the MAS promoter having a lower level of expression than p35S, populations of transformants created with this promoter show normally distributed expression levels (Perez-Gonzalez and Caro 2019). Thus, the MAS promoter can be used for functional screening of positive transformants in both of our shoot re-generation and rooting medium supplemented with bialaphos.

It is noteworthy that, before the initiation of stable transformation, we evaluated the effectiveness of gRNAs by using a convenient *Agrobacterium*-mediated transient assay method (Wang et al, the company manuscript submitted for review). As soybean plants have a long-life cycle (4-6 months), the estimation of the gRNAs’ effectiveness helps to avoid the waste of time and enhance the possibility of obtaining authentic gene-edited plants.

Although the soybean cultivars with natural variations on either *KTI1* (PI 68679) or *KTI3* (PI 542044) have been discovered, conventional breeding to develop new cultivars stacking with two mutant alleles via crossing will take a long time. In addition, linkage drag might lead to interference with the functions of genes located at the flanking sequences of mutant *kti1* or *kti3*. The limited genetic background of natural *kti1* and *kti3* mutants may also reduce the genetic diversity of soybean breeding lines with low TI trait, and it can be difficult to stack low TI trait with a bundle of various, desirable traits. In the present study, the *kti1/3* mutant line was created using cv. *WM82*, which has a genetic background distinct from accessions that harbor natural *kti1* and/or *kti3* mutations. Therefore, it offers a new recourse for breeding low TI traits in soybean practice.

Current soybean transformation protocol is genotype dependent, and only a few cultivars (*WM82, Jack, Thorne*, etc.) can be efficiently transformed (Yamada, Takagi et al. 2012). A mutant allele must be created in those transformable cultivars and bred into other elite cultivars via marker-assisted selection (MAS). Thus, creating a mutant allele tagged with convenient molecular markers is essential for MAS (Hasan, Choudhary et al. 2021). CRIPSR/Cas9-based genome editing can introduce small deletions/insertions to targeted genes, enabling us to develop molecular markers based on the sequences of the insertion and deletion mutation regions. In this study, we tested using single sgRNA and two sgRNAs for generation of mutagenesis on *KTI1* and *KTI3*, respectively (Figure 3B). Interestingly, we observed that all genotyped mutant lines carried an identical gene editing pattern of the *kti1* gene, where 66 nucleotides between the two gRNAs were deleted. The homozygous *kti1* allele can be identified in all tested seeds of the T0 generation, while homozygous *kti3* alleles were identified in some of those genotyped T0 seeds (Figure 4). Therefore, it is possible that two sgRNAs are more efficient for triggering the gene editing events in early generations of transgenic plants.

MAS has been widely implemented in plant breeding including soybean programs (Hasan, Choudhary et al. 2021). The selection marker of *kti3* has been developed based on its natural mutant allele, but the molecular marker for the natural mutation of *kti1* in PI 68679 is still not available (Gillman, Kim et al. 2015). Therefore, it is challenging to breed the natural *kti1*/*3* mutant alleles into a new cultivar via MAS. In this study, we created co-dominant markers that can distinguish between the wild and mutant alleles of *KTI1*/*KTI3* and *kti1*/*kti3* based on small deletions created by CRISPR/Cas9 machinery (Figure 7). In addition, a simple gel-electrophoresis-free method can be used to identify plants carrying mutant *kti1* and *kti3* alleles (Hirotsu, Murakami et al. 2010). Thus, MAS makes it possible to effectively breed the new mutant *kti1*/*3* alleles into other elite cultivars. Taken together, the whole experimental design may serve as a practical example of how to create and select mutant alleles in crop plants in the future.

## Conclusions

The present study developed non-transgenic, low TI soybean mutant in cv. William 82. The mutant gene alleles are tagged with convenient molecular markers that are suitable for high throughput marker-aided selection. We expect the low TI soybean mutant will be widely used to breed low-TI or TI-free soybean cultivars for commercial production in value-added meal industry and for stacking with other valuable agronomic traits in the future.

## Acknowledgements

This work was supported by the Virginia Soybean Board (#467059) and a seed grant from the Translational Plant Science Center at Virginia Tech. An integrated internal competitive grant from the College of Agriculture and Life Sciences at Virginia Tech and Virginia Agricultural Experiment Station (VA160144).

## Author contributions

B.Y.Z. and Z.W. designed the experiments and analyzed data. Z.W. performed experiments, analyzed data, and wrote the manuscript. Z.S., L.R., C.S., J.L., and P.B. also performed experiments. B.Y.Z., B.Z., Z.S., L.R., and P.B. edited the manuscript. B. Z. supervised the project.

## Methods

### Plant materials and growth conditions

Soybean plants were grown in 2.5-gallon pots using Miracle-Gro all-purpose potting soil mix in Keck Greenhouse at Virginia Tech (14h/10h light/dark cycle at 25 °C/20 °C) for the experiments described herein. The plants were watered by an automatic irrigation system. Soybean transformation was performed at the plant transformation facility at Iowa State University as previously described (Paz, Martinez et al. 2006, Luth, Warnberg et al. 2015, Ge, Yu et al. 2016). The plant growth indicators and maturity period days of *WM82* and progeny plants of the T1 generation derived from lines #2 and #5 were measured in the green house. The 4-week-old T1 soybean plants were used for genotyping. The seeds of T1 plants were used for seed weight analysis.

### Constructing a soybean KTI gene map

The gene map showing locations of *KTI* genes on soybean chromosomes was made using MapInspect. Locations of all *KTI* genes were obtained from the Phytozome database and plotted on their respective chromosomes.

### Bacterial growth

*E. coli* strains *DH5α* and *C41* (DE3) (Lucigen, Middleton, WI) were grown on Luria agar medium at 37 ℃. *Agrobacterium tumefaciens* (*A. tumefaciens*) *EHA105* was grown on Luria agar medium at 28 ℃ (Zhao, Dahlbeck et al. 2011). *E. coli* antibiotic selections used in this study were as follows: 50 μg/ml kanamycin, 100 μg/ml carbenicillin, 100 μg/ml spectinomycin. *A. tumefaciens* antibiotic selection were 100 μg/ml rifampicin, and/or 100 μg/ml spectinomycin.

### Cloning

The open reading frames (ORFs) of *KTI1* and *KTI3*, were amplified from the genomic DNA of *WM82*. The *KTI1*_Δ66bp_, truncated ORF of *KTI1*, was amplified from the genomic DNA of mutant soybean plant #2-1. All PCR primers with annotations are listed in Table S1. The genes/fragments were then cloned into a pDonr207 plasmid (Thermo Fisher Scientific) for future use.

T1 plant genomic DNA (gDNA) was used as the templates to amplify KTI1 and/or its mutant allele, and KTI3 and/or its mutant allele by PCR. The purified PCR fragments were used for genotyping by Sanger sequencing at Virginia Tech Genomic Sequencing Center and cloned to the PCR8/GW/TOPO vector by TA cloning (Invitrogen) for molecular marker tests.

In order to apply the CRISPR/Cas9 system to gene editing in soybean, we modified our current CRISPR/Cas9 construct (Liu, Miao et al. 2016). The cassette consists of a MAS promoter, the bialaphos resistant gene, and a MAS terminator that was amplified using plasmid DNA of pEarleyGate101 as the template. All PCR primers with annotations are listed in Table S1. The cassette was assembled to the backbone of CRISPR/Cas9 construct using Gibson Assembly® Cloning Kit (New England Biolabs Inc). The gRNAs targeting*KTI1* and *KTI3* were synthesized in one cassette at GenScript Biotech Corp. The backbone of the new CRISPR/Cas9 construct and the fragment of gRNAs were assembled together using Gibson Assembly® Cloning Kit.

### Expression analysis of KTI Genes in WM82

RNA sequencing data, in FPKM (fragments per kilobase of transcript per million fragments mapped), of 38 *KTI* genes in 31 different tissue types from Williams 82 were acquired through the Gene Networks in Seed Development database (http://seedgenenetwork.net/sequence). Construction of the heatmap to visualize expression data was done using the heatmap.2 function from the ggplot2 package in R. A green/blue color gradient was chosen to show expression with blue representing little to no expression and green representing high expression. The code for the heatmap is as follows: *heatmap*.*2(x=KTI Expression, main = “KTI Expression In Different Soybean Tissue”, notecol=“black”, density*.*info=“none”, trace=“none”, margins =c(12,9), col=my_palette, breaks=col_breaks, dendrogram=“row”,Colv=“NA”, ylab= “Genes”, xlab= “Tissue Type”, cexCol=*.*9,cexRow =* .*8)*

### RNA isolation and real-time PCR

All RNA was extracted from V98-9005 and V03-5903 seeds using TRIzol reagent (Thermo Fisher Scientific) according to the manufacturer’s instructions. Any DNA residue was eliminated by treating with UltraPure DNase I (Thermo Fisher Scientific). The integrity and quantity of total RNA were determined by electrophoresis in 1% agarose gel and a NanoDrop ND-1000 spectrophotometer (NanoDrop Technologies, Wilmington, DE). cDNA synthesis was performed using the SuperScript III First-Strand RT-PCR Kit (Thermo Fisher Scientific) with an oligo-dT primer based on the manufacturer’s instructions. Real-time PCR was conducted with cDNA as the template using the Quantitect SYBR Green PCR kit (Qiagen) according to the manufacturer’s protocol. Oligo primers are listed in Table S1. The soybean *ELF1B* gene was used as reference gene, and data is presented as ΔCT (Jian, Liu et al. 2008).

### Expression and purification of KTI1, KTI1_Δ22aa_, and KTI3 proteins

The *KTI1, KTI1*_Δ66bp_, and *KTI3* genes in pDonr207 were subcloned into a Gateway compatible pET28a destination vector via a LR® Gateway cloning kit (Thermo Fisher Scientific) (Earley, Haag et al. 2006, Liu, Miao et al. 2016). The plasmids were transformed into *E. coli C41* cells (Lucigen). KTI1, KTI1_Δ22aa_, and KTI3 proteins were expressed and purified following a procedure as previously described [43]. Protein purity was evaluated by SDS-PAGE. The protein concentration was determined by a protein assay kit (Bio-Rad) using bovine serum albumin as standard (Han, Zhou et al. 2015).

### Standard bioassay to measure trypsin inhibitor activity

A TI activity bioassay was performed following American Association of Cereal Chemists Official Method 22-40 (AACC, 1999) with some modifications previously reported by Rosso et al.(2018). Briefly, 30 mg of finely ground soybean seed powder was mixed with 3 mL of 9 mM HCl (pH 2.0). The mixture was shaken for 1 h at room temperature. 2 mL of the extracts was centrifuged at 10,350 rpm for 20 min at room temperature, and the supernatant was diluted by 10 times with 9 mM HCl for measuring TI activity. A TI activity assay was performed in a 96-well plate format following the same steps described in Rosso et al. (2018). Each sample row was repeated three times. Portions of diluted HCl extracts (0, 20, 30, 40, and 60 µL) or 50 µg recombinant proteins of KTI1, KTI1_Δ22aa_, and KTI3 were pipetted into the microplate wells, and the volume was adjusted to 60 µL with 9 mM HCl. 60 µL of extractant was used as a sample blank and 60 µL of water were used as a substrate blank. 60 µL of trypsin (from bovine pancreas, Sigma-Aldrich T8003) solution was added to each sample well, and the microplates were placed in an oven at 37°C for 15 min. After the incubation, 150 µL of BAPNA substrate pre-warmed at 37°C was added to all wells, and the plates were incubated for exactly 10 min at 37°C. The reaction was stopped by adding 30 µL of acetic acid solution to all wells. The absorbance of each well was read on a plate reader (FLOUstar Omega, BMG Labtech) at 410 nm for 30 s after shaking at 700 rpm.

### HPLC method to quantify kunitz trypsin inhibitor

The HPLC method to quantify *KTI* was performed following the method developed by Rosso et al. (2018). Briefly, 10 mg of finely ground soybean seed powder was mixed with 1.5 mL of 0.1 M sodium acetate buffer (pH 4.5). Samples were vortexed and shaken for 1 h at room temperature. The sample was centrifuged at 12,000 rpm for 15 min. 1mL of the supernatant was filtered through a syringe with an IC Millex-LG 13-mm mounted 0.2-mm low protein binding hydrophilic millipore (polytetrafluoroethylene [PTFE]) membrane filter (Millipore Ireland). The *KTI* in solution was separated on an Agilent 1260 Infinity series (Agilent Technologies) equipped with a guard column (4.6 × 5 mm) packed with POROS R2 10-mm Self Pack Media and a Poros R2/H perfusion analytical column (2.1 × 100 mm, 10 um). The mobile Phase A consisted of 0.01% (v/v) trifluoroacetic acid in Milli-Q water, and the mobile Phase B was 0.085% (v/v) trifluoroacetic acid in acetonitrile. The injection volume was 10 mL and the detection wavelength was 220 nm.

### Development of molecular selection markers with a gel electrophoresis free method for high throughput screening

The transgene free and double homozygous mutant line, #5-26, was selected for the development of molecular selection markers. Based on the genotyping data of #5-26, two pairs of markers were designed: ZW1 with ZW2 or ZW3. ZW1 is the common reverse primer for both *KTI1* and *kti1*, while ZW2 and ZW3 are two reverse primers matched with unique sequences in *KTI1* and *kti1*, respectively. Similarly, two pairs of molecular markers, ZW4 with ZW5 or ZW6, were designed. ZW4 is the common reverse primer for both *KTI3* and *kti3*, while ZW5 and ZW6 are two reverse primers matched with unique sequences in *KTI3* and *kti3*, respectively. The gDNA of *WM82*, #5-26 (homozygous mutants of both *kti1* and *kti3*), #5-9 (homozygous mutant of *kti1* while heterozygous mutant of *kti3*) and #2-30 (homozygous mutant of *kti3* while heterozygous mutant of *kti1*) were used as templates to test the efficiency and reliability of these markers in PCR.

PCR amplifications were performed in a total volume of 20 μl containing 50 ng of gDNA, 0.5 μM each of forward and reverse primers (Table S1), 10 μl 2X BioMix Red (Bioline) and ddH_2_O. The PCR program was set to be 95 °C for 5 min for pre-denature, followed by 35 cycles of denaturation at 95 °C for 30 s, annealing at 55 °C for 30 s, extension at 72 °C for 30 s, followed by final extension at 72 °C for 5 min.

In order to screen the large-scale progenies derived from crosses between soybean elite cultivars carrying wild type *KTI1*/*3* and the newly developed mutant plant carrying *kti1*/*3*, a simple gel electrophoresis-free method was designed. 1X sybrgreen dye (Thermo Fisher) was added to complete PCR reactions, and the solution was incubated for 10 mins at 75°C before placing in the gel doc (Biorad) for imaging the fluorescent signals. Only the positive PCR products with the dye will display fluorescent signals while the failed PCR will not show signals.

### Statistical data analysis

Analytical experiments were performed with at least three technical replicates. Statistical significance was based on one-way ANOVA test for multiple comparisons. Data was analyzed using JMP Pro14. Values of P<0.05 were considered significant.

## Figure Legends

**Figure S1.**
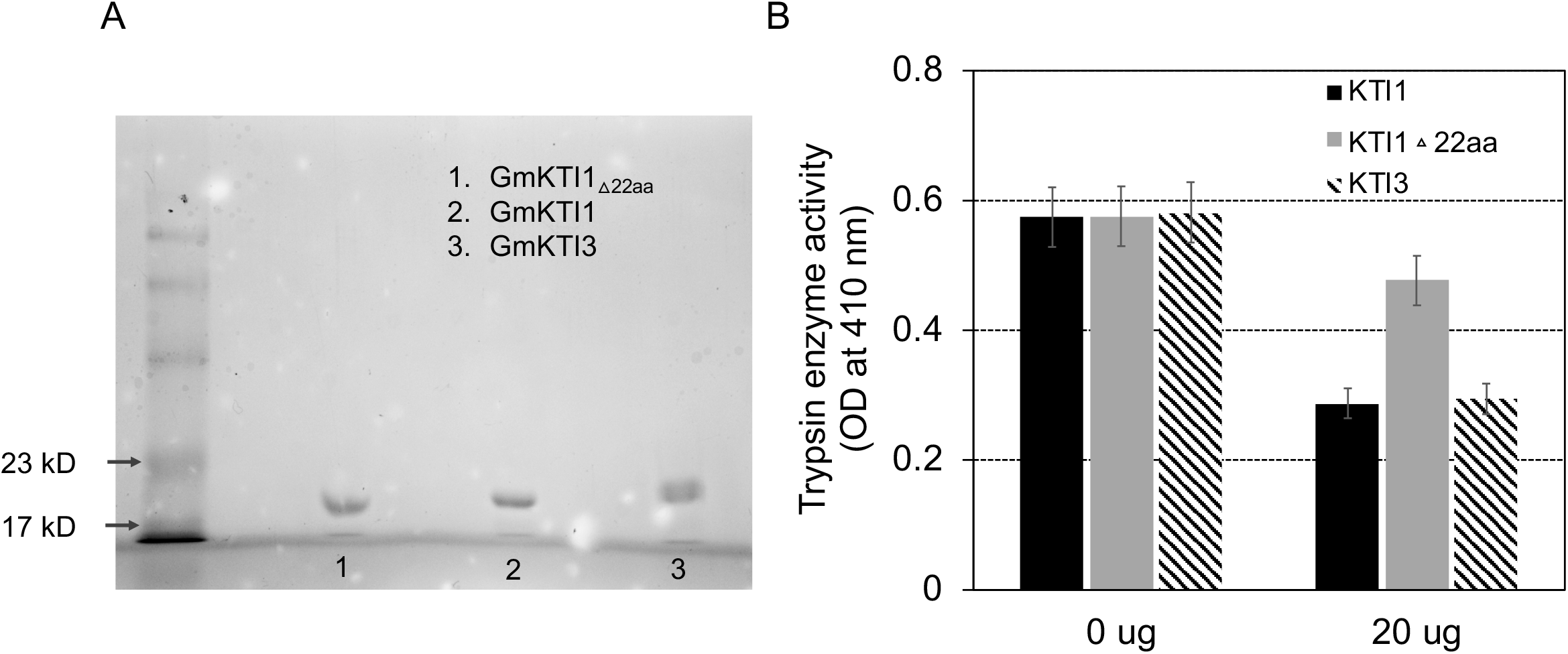
SDS-PAGE of purified recombinant proteins and in-frame mutated protein of KTI1_Δ22aa_ nearly lost the TIA. (A) SDS-PAGE was used to assess the purity of three recombinant proteins, KTI1, KTI3, and KTI1_Δ22aa_. (B) Purified proteins of KTI1 and KTI3, but not KTI1_Δ22aa_ were able to inhibit the trypsin activity *in vivo*. Experiments were conducted with three technical replicates and obtained similar results.

**Table S1.**
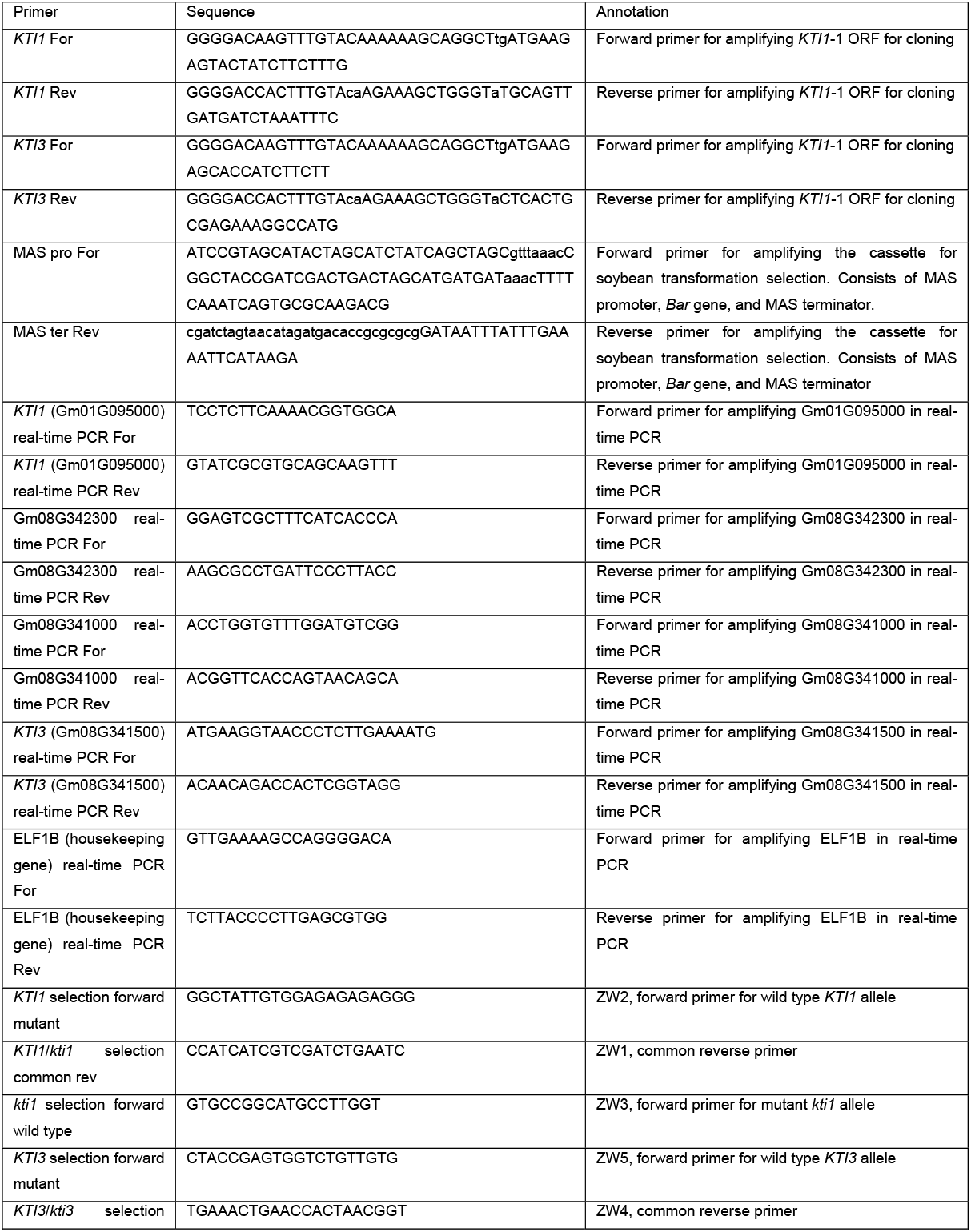
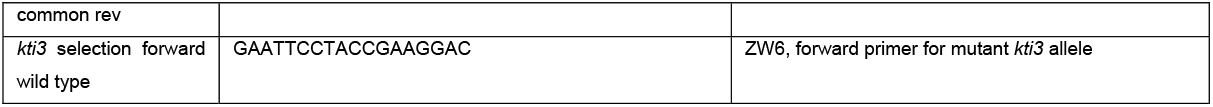
Oligo primers used in this study.

